# Long-term consequences of fostering: Single egg fostering leads to decreased survival in zebra finch females, but not in males

**DOI:** 10.64898/2026.02.01.703085

**Authors:** Barbara A. Caspers, Sabine Kraus, Sarah Golüke, Marta Rossi

## Abstract

Cross-Fostering, i.e., the exchange of eggs or hatchlings, is a widely used technique, to disentangle genetic from environmental effects or to manipulate the clutch size. In most bird species, this manipulation is easily accepted by the social parents, leading to the conclusion that fostering has no detrimental effect. Using a dataset of four cohorts (N=298) of zebra finches (*Taeniopygia castanotis*), in which we fostered routinely a single egg into another nest of zebra finches, we explored potential short- and long-time effects of fostering. Noteworthy, these experiments were not designed to test this hypothesis. The objective of the egg fostering experiments was to test for parental recognition (Caspers et al. 2017) and mate choice decisions (Golüke 2018). Consequently, the aim of the present study is purely explorative. Our study confirmed previous findings that fostering has no short-term effects on the morphology and growth rates of the chicks, neither in males nor in females. However, we found that fostering has a sex-specific long-term effect. Females originating from fostered eggs had a significantly reduced lifespan compared to those from non-fostered eggs. Conversely, the lifespan of fostered males was similar to that of non-fostered males. All birds were housed in large groups, experiencing the same conditions after nutritional independence (day 35). Therefore, we can only speculate that fostering might result in early developmental stress, which may affect the individual fitness of females later in life, ultimately leading to shorter lifespans.

## Introduction & Results

Cross-fostering, i.e. the exchange of eggs or hatchlings, allows to experimentally disentangle genetic components from environmental factors, to manipulate brood sizes and to investigate the effect of non-genetic inheritance on various life history. This manipulation is widely used in almost all aspects of studies on bird biology, from kin recognition (Sharp et al. 2004) to sibling competition (Lattore et al. 2019), as well as studies on parental care and conflict (Royle et al. 2002, Wright & Cuthill 1990), and mating preferences (Clayton, 1990). Parents typically readily accept new members into the family and feed them as if they were their own, indicating no short-term effects of cross-fostering.

Given our aim to test the impact of the nest environment on parent-recognition (Caspers et al. 2017) and mating preference (Golüke 2018), we used a standardised cross-fostering design, in which we fostered single eggs into the nest of conspecifics. From 2015 to 2018, we produced a total of 298 chicks (148 females, 150 males) across 4 cohorts of zebra finches, comprising 115 non-foster females, 33 foster females, 112 non-foster males, and 38 foster males. We followed each individual during the nestling period until nutritional independence (35 days) and measured the weight at the day of hatching (day 0), day 3, day 6, day 10, and day 35. As suggested by other studies, we found that the body weight gain was neither affected by brood size, nor by foster status or sex (Table 1, Figure 1). Body weight increased over time with a similar rate in female and male chicks, either fostered or non-fostered, suggesting that under our conditions cross-fostering did not have an apparent effect in the short-term (Table 1, Figure 1).

**Table 1.**
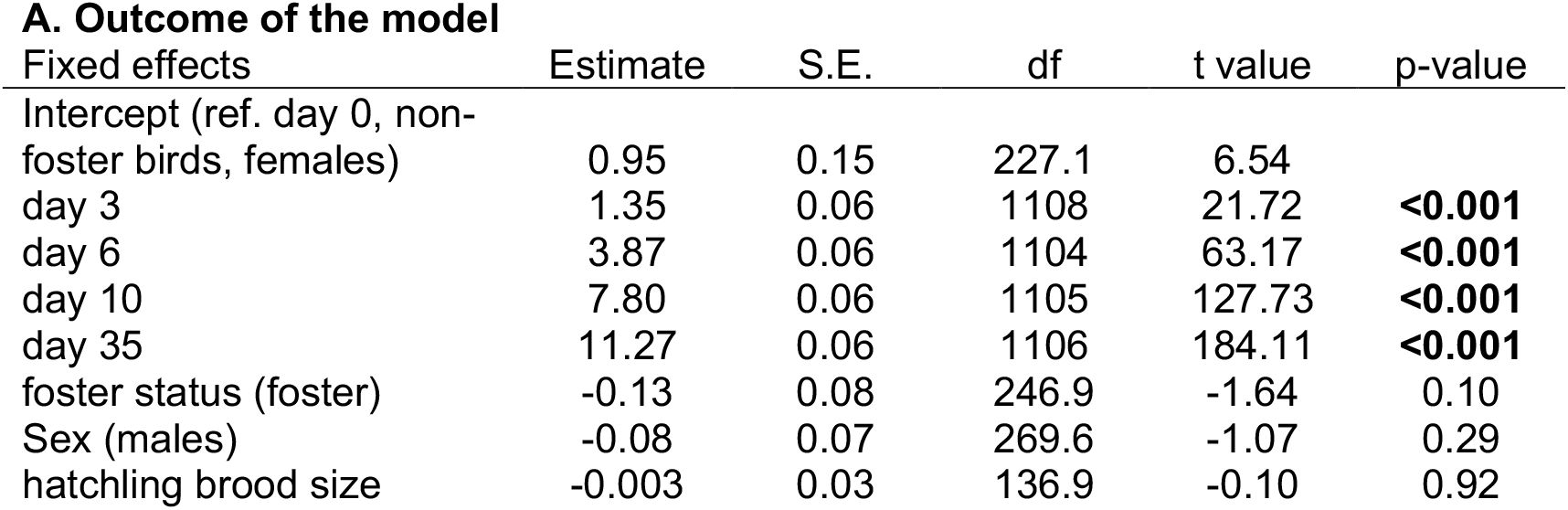
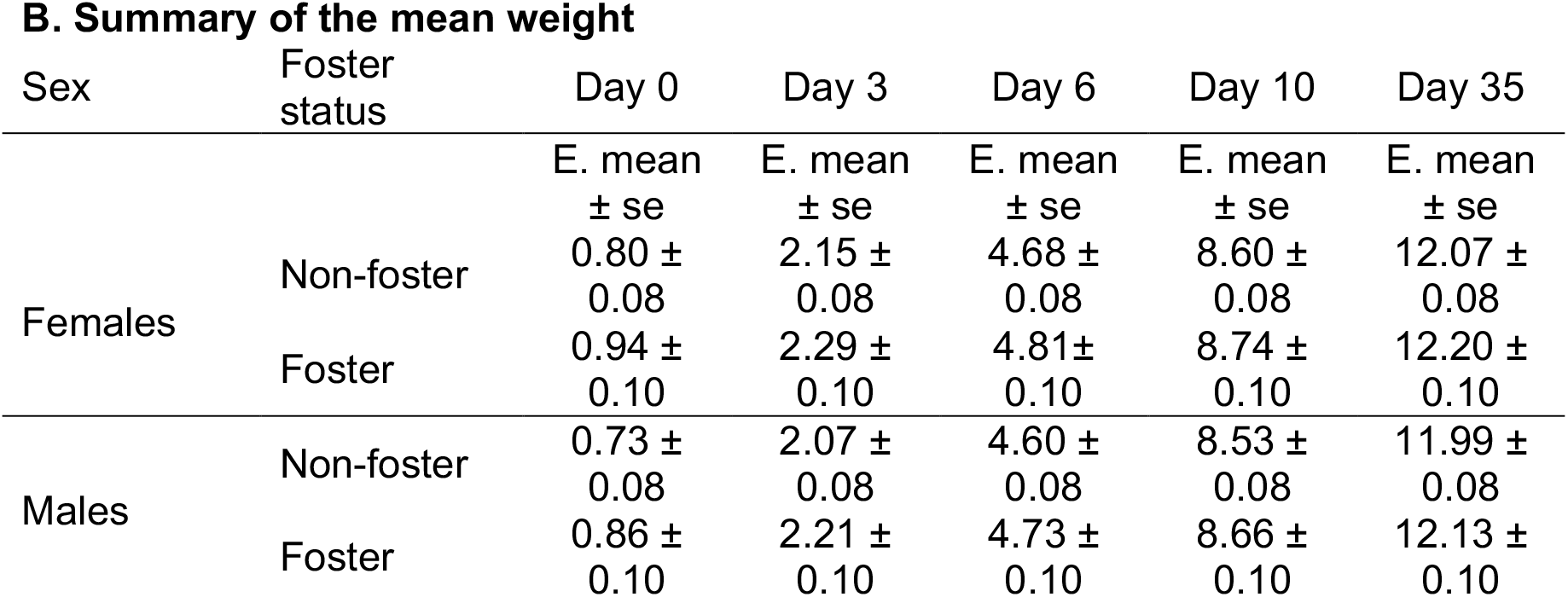
The weight increase of foster chicks was similar to that of non-fostered chicks and did not differ between sexes. **A)** Outcomes of the LME models investigating the effect of foster status (foster vs. non-foster), sex (females vs. males) and hatchling brood size (continuous; brood size range between 1 and 7 chicks) on the weight increase of zebra finch chicks in early life (day 0, 3, 6, 10, and 35). **B)** Summary of the mean weight of non-foster and foster chicks at each time point. “lmer (chick weight ∼ days + foster status + sex + hatchling brood size + (1|ID) + (1|mother)”

**Figure 1.**
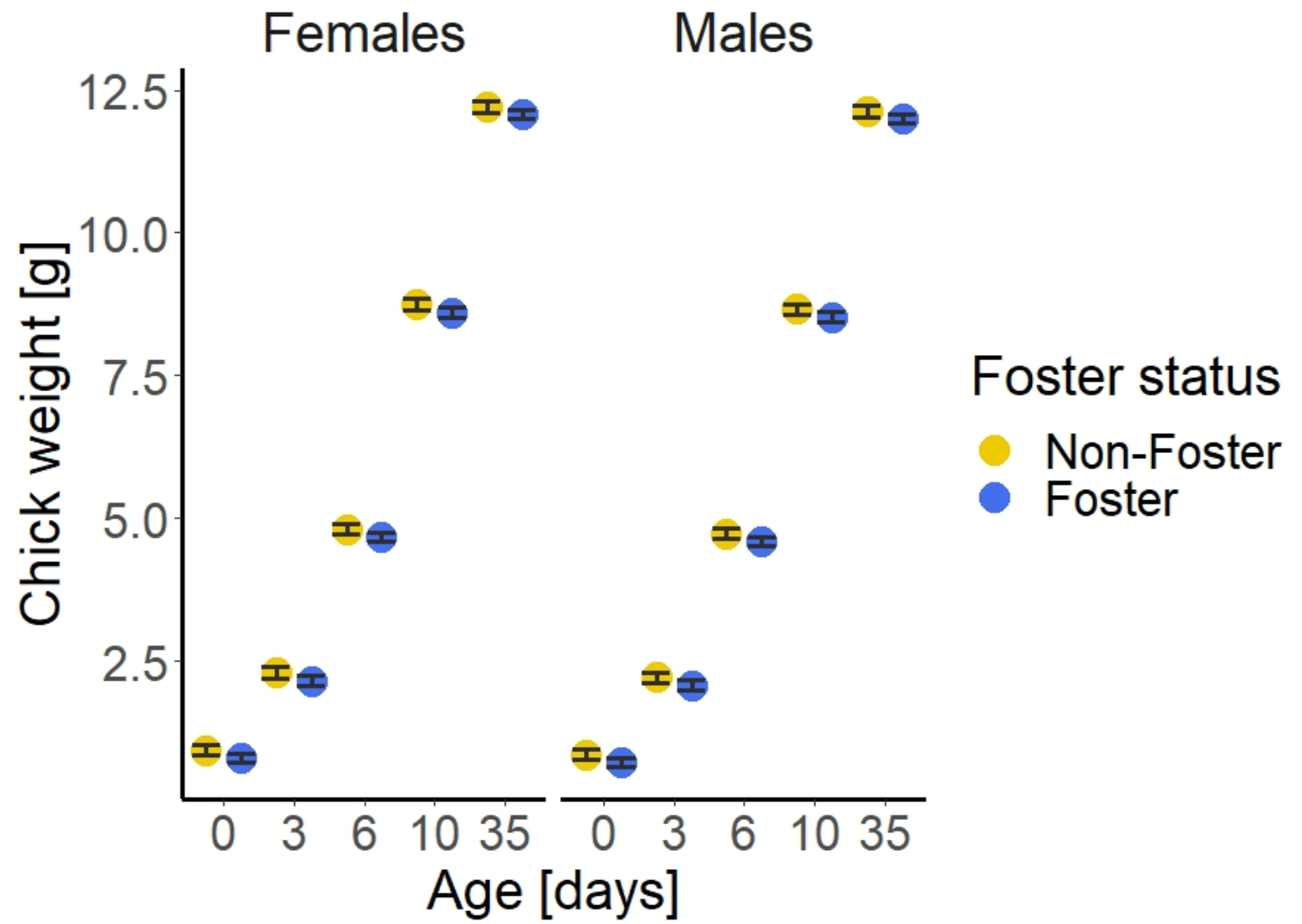
The weight increase of foster chicks was similar to that of non-fostered chicks and did not differ between sexes. Circles represent means estimated fitted from LMEM, error bars represent the standard errors.

After nutritional independence (day 35) all chicks were moved to mixed group aviaries, consisting of 20-50 birds, depending on the size of the aviary, in which a minimum of one adult pair was present. Each housing room consisted of two to four aviaries. All birds remained at the department until their death, which was noted on the aviary list. In addition, during the yearly inventory, all birds of our aviaries were captured.

We performed a survival analysis to exploratively investigate potential long-term effects of cross fostering and found an unexpected sex-specific long-term effect of egg fostering. Female birds originating from fostered eggs showed a significantly decreased probability of survival compared to those that were not fostered (χ^2^ = 5, df = 1, p = 0.03), while there was no difference in survival between fostered and non-fostered male birds (χ^2^ = 0, df = 1; p = 0.8; Table 2, Figure 2, Table S1,S2). In addition, we investigated the effect of breeding experience on the survival probability of female birds that reached sexual maturity (lifespan >100 days). The survival of virgin females was similar to those that had bred, for fostered birds (Log-rank test; 21 virgins, 12 bred: χ^2^ = 1.8, df = 1, p = 0.2). However, the survival of virgin females was lower than for those that had bred in the non-foster birds (Log-rank test; 72 virgins, 43 bred: χ^2^ = 4.5, df = 1, p = 0.03).

**Table 2.** Log-rank test to compare survival probability of foster and non-foster male birds.

**A) females**
| Foster status | N | N Observed | Expected | (O-E)<sup>2</sup>/E | (O-E)<sup>2</sup>/V |
| --- | --- | --- | --- | --- | --- |
| Non-foster birds | 115 | 45 | 51.5 | 0.82 | 4.96 |
| Foster birds | 33 | 18 | 11.5 | 3.69 | 4.96 |
| X<sup>2</sup>= 5 on 1 degrees of freedom, **p=0.03** |  |  |  |  |  |
| **B) males** |  |  |  |  |  |
| Foster status | N | N Observed | Expected | (O-E)<sup>2</sup>/E | (O-E)<sup>2</sup>/V |
| Non-foster birds | 112 | 31 | 31.5 | 0.009 | 0.04 |
| Foster birds | 38 | 12 | 11.5 | 0.03 | 0.04 |
| X<sup>2</sup>= 0 on 1 degrees of freedom, p=0.8 |  |  |  |  |  |

**Figure 2.**
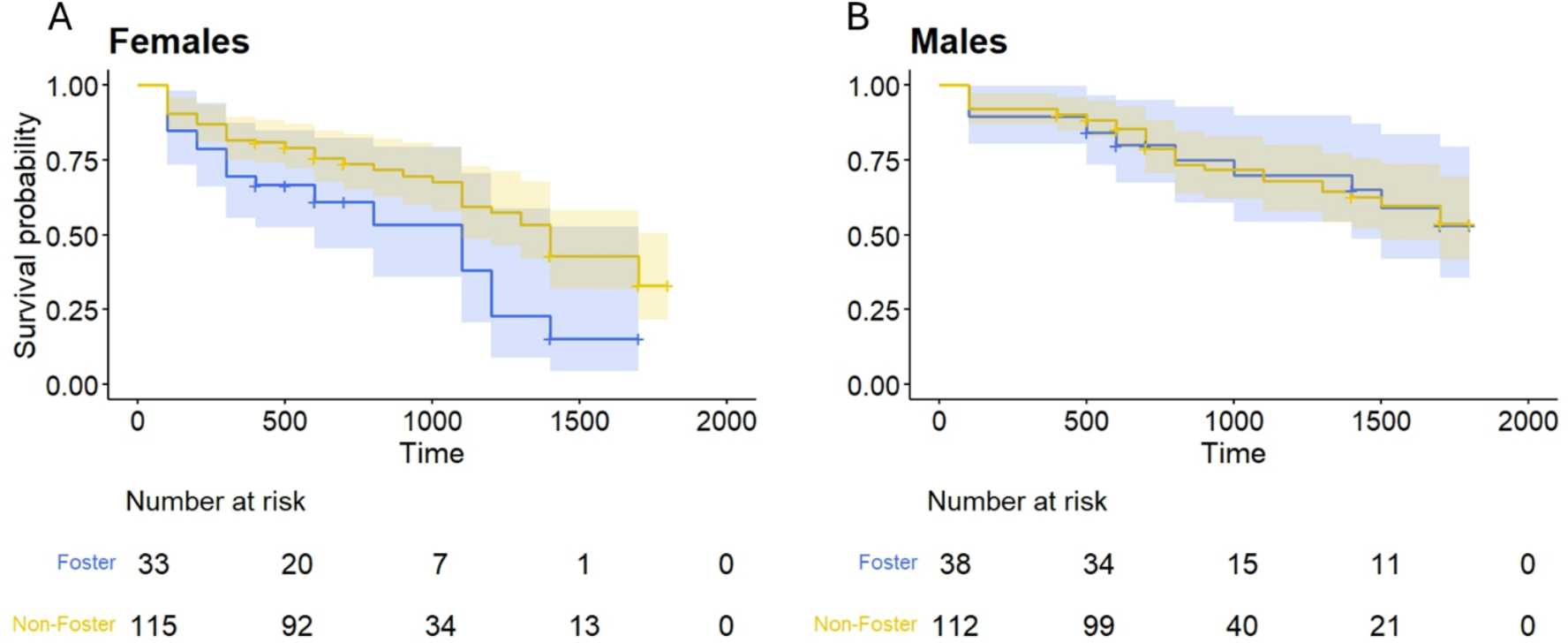
The probability of survival was lower for female birds raised from fostered eggs compared to non-fostered ones, while it was similar between fostered and non-fostered male birds. Vertical tick-marks indicate censored birds.

## Discussion

Cross-fostering is a widely used method and as suggested, we did not find any evidence for a short-term effect of fostering, indicating that adult birds do not preferentially feed their own chicks and do not discriminate against kin (Beecher 1991, Kempenaers & Sheldon 1996, but see Golüke et al. 2021). However, this explorative study uncovered a robust sex-specific effect of fostering on the survival probability, with females originating from fostered eggs having a significantly reduced life expectancy. Furthermore, our data suggests that longevity is also influenced by the possibility to reproduce, with virgin females having a lower life expectancy that females that bred.

Life expectancy varies considerably between and among species, and sex is thought to have a large effect on longevity (Xirocostas et al. 2020), with the heterogametic sex having a reduced life expectancy (i.e. female in birds). Accordingly, we showed that female zebra finches had a lower survival probability than males. In addition, several cellular mechanisms such as IGF-1 signalling, telomere shortening, oxidative stress, and stress resistance seem to play an important role in controlling the rate of aging (Monaghan 2014; Monoghan et al. 2012; von Zglinicki et al., 2001, Viblanc et al. 2020). Environmental conditions, such as nutrition, are also associated with aging (Fontana et al. 2010). However, the intertwined relationship between external factors, cellular mechanisms and individuals’ longevity remains poorly understood.

Sexual activity and the possibility to reproduce influenced life expectancy in non-fostered zebra finch females. This is quite unexpected, as an investment in reproduction is costly and supposed to decrease longevity (Brooks & Garratt 2016). However, a similar finding has been described in the Ansell’s mole-rat *Cryptomys anselli*, in which reproductively active individuals had a significantly higher life expectancy than those that were not allowed to breed (Dammann and Burda 2006). Due to the purely descriptive and explorative nature of our study we cannot rule out the possibility that breeding females were in overall better bod condition that virgin females, and were therefore selected to breed, but we think this is highly unlikely as we choose breeding pairs randomly and not based on body condition. Like what have been found in the Ansell’s mole rat (Dammann and Burda 2006), our results might show a general positive effect of reproduction on longevity, at least in captive bred animals.

Although fostering did not show an apparent short-term phenotypic consequence in early life, the most plausible explanation for the differences in survival probability lies in the early environment, namely the lack of close relatives for fostered individuals. Indeed, after nutritional independence (day 35) all birds were exposed to the same environmental conditions, and food was provided *ad libitum*. Therefore, we suggest that the nestling period was the only window of time in which differences between foster and non-foster chicks may have arisen, potentially leading to increased stress, either during early development or later. In birds, increasing evidence suggests that early developmental stress such as food scarcity, sibling competition, and predation could have long-term consequences on life expectancy, even if there are no obvious consequences at the time of the stressful event (Metcalfe and Monaghan, 2001). Furthermore, male and female birds show different responses to early developmental stress (López-Arrabé et al. 2016). To cope with environmental stressors and maximise the chance of survival, animals can modulate their physiological response by increasing the secretion of stress hormones such as corticosterone and testosterone. Increased glucocorticoid levels may be associated with increased oxidative stress and telomere shortening, and both processes seem to be linked with longevity (Costantini et al., 2008; Haussmann and Marchetto, 2010; Monaghan et al., 2009; Pegan et al., 2019). A correlation between increased physiological stress and telomere shortening rates has been shown in European shags (Herborn et al., 2014), and in zebra finches the telomere length in early life (25 days) correlates with future lifespan expectancy (Heidinger et al., 2012).

The explorative nature of our study allows us, unfortunately, only to speculate about the potential mechanisms. While it has been shown that parents do not discriminate against unrelated chicks (Kempenaers & Sheldon 1996), it is plausible that, according to kin selection theory (Hamilton 1964), sibling competition might increase in broods with a foster chick, particularly resulting in heightened competition for the fostered chick (Briskie et al. 1994, Boncoraglio et al. 2006). Similar findings have been found in mammals, e.g. piglets, in which being raised in mixed litters had detrimental effects on heavy piglets, but beneficial effects on light piglets (Vande Pol et al. 2021). In wild cavies, cross fostered individuals were less bold, grew slower and showed elevated resting metabolic rates (Kraus et al. 2020) and in mice a sex specific effect of cross fostering has been found (Bartolomucci et al. 2001).

Zebra finch nestlings have been shown to be able to discriminate kin and modulate their behaviour based on the ratio of related and unrelated nest mates (Krause et al. 2012). Moreover, zebra finch chicks can discriminate between the odour of their biological parents and that of an unfamiliar parent at the day of hatching (Caspers et al. 2017) and retain a preference for the odour of their biological mother even after egg cross-fostering (Caspers et al. 2017). Blue tit chicks discriminate between the odour of a familiar related chick and an unfamiliar unrelated chick (Rossi et al. 2017), but do not show any sign of discrimination when related and unrelated nestmates are equally familiar (Schlatmann et al. 2025). Although not tested in our study, zebra finch chicks may be able to discriminate between related and unrelated nestmates. A single foster chick surrounded by a majority of unrelated nestmates may suffer from either by direct mobbing or increase the duration and intensity of its begging to compete against them.

It remains unclear why fostering had a sex-specific long-term effect, impacting exclusively the longevity of females. A partial explanation could be that females are more susceptible to stressful circumstances than males. Being the heterogametic sex, females may express undesirable physiological and morphological characteristics through the expression of recessive mutations more likely than males (Xirocostas et al. 2020). Further ad hoc experiments and investigation during the nestling phase such as quantifying behavioural differences between foster and non-foster chicks, hormonal stress assay, and telomere length and attrition measurements are needed to understand the underpinning mechanisms and the reasons why female foster birds appear to be more susceptible than foster males. Nevertheless, we showed that egg fostering had a strong impact on the survival probability of females, even in the absence of developmental short-term effects. Such effects should be considered when performing cross-fostering studies.

## Methods

### Beeding, housing and cross-fostering procedure

Pairs of birds from our stock of domesticated zebra finches, *Taeniopygia castanotis* (Griffith et al. 2017, note that the species name was changed from *T. guttata* to *T. castanotis*) were randomly chosen and allowed to breed in two-compartment breeding cages (W: 80 cm, H: 30 cm, L: 40 cm, JAKO). We conducted daily nest checks, recording the onset of nest building, egg laying, the number of eggs in the nest, the number of chicks hatched, and the number of chicks in the nest. When two cages started laying eggs simultaneously, we cross-fostered the third egg between the two nests on the same day the egg was laid. We always fostered the third egg to avoid any influence of laying order and potential differences in begging intensity that might arise from it (Gilby et al. 2011). In case three nests were available at the same time, we circularly exchanged the third egg between the three nests (i.e., egg from nest A into nest B, egg from nest B into nest C, and egg from nest C into nest A).

All eggs, including the non-fostered eggs were marked with a permanent marker at the day of egg laying. This way we were able to identify the fostered chick at hatching, as usually only one chick per day hatches. In case two chicks hatched on the same day, we performed paternity analysis (Caspers et al. 2017) to determine the genetic parents of the chicks. Shortly after hatching, all chicks were marked individually by cutting the down feathers in a distinctive pattern (Adam et al. 2014). On day 10, chicks were individually marked with numbered grey plastic rings.

### Statistical Analysis

All statistical analyses were conducted in R version 4.3.0 (R Core Team 2023) in R-Studio (version 2025.9.1.401 R Development RStudio Team 2020). To create the graphs, we used the packages ggplot2 (Wickham 2016) and survminer (Kassambara et al. 2025). We fitted LME models by restricted maximum likelihood estimation (lmer function, package ‘lme4’) (Bates et al., 2014). Significances of the fixed effects were determined using Satterthwaite’s method for estimation of degrees of freedom (package ‘lmerTest’) (Kuznetsova et al., 2017) and non-significant interactions (P>0.05) were removed. Estimated marginal means and pairwise comparisons were obtained using the ‘lsmeans’ package (Lenth and Lenth, 2018).

We used Kaplan-Meier survival analysis and log-rank test to determine the effect of cross-fostering on the probability of survival of female and male Zebra finch birds (‘survival’ package) (Therneau and Lumley, 2014). The Kaplan-Meier estimator is a non-parametric statistic that includes censored observations (i.e. incomplete observations), such as birds that had not died within the observation period.

## Supporting information

tables with estimates for the survival curves

## Supplementary material

Table S1 and Table S2

## Author contributions

BC and SG conceived the study, MR and SK analysed the dataset, SG collected the data, BC and MR wrote the manuscript.

## Acknowledgments

We thank all student helpers during data collection Jonas Tebbe, Anna-Kathrin Sonntag, Marina Inhofer, Madeleine Paul. We also thank Elke Hippauf for conducting the paternity analysis in cases where the foster status wasn’t clear. Furthermore, we thank Oliver Krüger for logistical support.

## Abbreviations

CI: coefficient interval
E: expected
LME: linear mixed effect
O: observed
se: standard error
V: variance

