## Supplementary material for "Long-term consequences of fostering: Single egg fostering leads to decreased survival in zebra finch females, but not in males": tables with estimates for the survival curves

**Table S1. Female foster birds showed a significantly lower survival probability compared with non-fostered birds.** Estimates of the survival curve using the Kaplan-Meier method for non-foster female birds (A), and foster female birds (B).

**A. Non-foster birds**

| time | n. risk | n. event | survival | se | lower 95% CI | upper 95% CI |
|---|---|---|---|---|---|---|
| 100 | 115 | 11 | 0.90 | 0.03 | 0.85 | 0.96 |
| 200 | 104 | 4 | 0.87 | 0.03 | 0.81 | 0.93 |
| 300 | 100 | 6 | 0.82 | 0.04 | 0.75 | 0.89 |
| 400 | 94 | 1 | 0.81 | 0.04 | 0.74 | 0.89 |
| 500 | 92 | 2 | 0.79 | 0.04 | 0.72 | 0.87 |
| 600 | 47 | 2 | 0.76 | 0.04 | 0.68 | 0.85 |
| 700 | 38 | 1 | 0.74 | 0.05 | 0.66 | 0.83 |
| 800 | 36 | 1 | 0.72 | 0.05 | 0.63 | 0.82 |
| 900 | 35 | 1 | 0.70 | 0.05 | 0.61 | 0.81 |
| 1000 | 34 | 1 | 0.68 | 0.05 | 0.58 | 0.79 |
| 1100 | 33 | 4 | 0.59 | 0.06 | 0.49 | 0.73 |
| 1200 | 29 | 1 | 0.57 | 0.06 | 0.46 | 0.71 |
| 1300 | 28 | 2 | 0.53 | 0.06 | 0.42 | 0.68 |
| 1400 | 26 | 5 | 0.43 | 0.07 | 0.32 | 0.58 |
| 1700 | 13 | 3 | 0.33 | 0.07 | 0.22 | 0.51 |

**B. Foster birds**

| time | n. risk | n. event | survival | se | lower 95% CI | upper 95% CI |
|---|---|---|---|---|---|---|
| 100 | 33 | 5 | 0.85 | 0.06 | 0.73 | 0.98 |
| 200 | 28 | 2 | 0.79 | 0.07 | 0.66 | 0.94 |
| 300 | 26 | 3 | 0.70 | 0.08 | 0.56 | 0.87 |
| 400 | 23 | 1 | 0.67 | 0.08 | 0.52 | 0.85 |
| 600 | 12 | 1 | 0.61 | 0.09 | 0.45 | 0.82 |
| 800 | 8 | 1 | 0.54 | 0.11 | 0.36 | 0.79 |
| 1100 | 7 | 2 | 0.38 | 0.12 | 0.21 | 0.71 |
| 1200 | 5 | 2 | 0.23 | 0.11 | 0.09 | 0.59 |
| 1400 | 3 | 1 | 0.15 | 0.10 | 0.04 | 0.53 |

**Table S2. Male foster birds show a survival probability similar to that of non-foster birds.** Estimates of the survival curve using the Kaplan-Meier method for non-foster male birds (A), and foster male birds (B).

**A. Non-foster birds**

| time | n. risk | n. event | survival | se | lower 95% CI | upper 95% CI |
|---|---|---|---|---|---|---|
| 100 | 112 | 9 | 0.92 | 0.03 | 0.87 | 0.97 |
| 400 | 103 | 2 | 0.90 | 0.03 | 0.85 | 0.96 |
| 500 | 99 | 2 | 0.88 | 0.03 | 0.83 | 0.95 |
| 600 | 60 | 2 | 0.85 | 0.04 | 0.79 | 0.93 |
| 700 | 52 | 4 | 0.79 | 0.05 | 0.70 | 0.88 |
| 800 | 44 | 3 | 0.74 | 0.05 | 0.64 | 0.84 |
| 900 | 41 | 1 | 0.72 | 0.05 | 0.62 | 0.83 |
| 1100 | 40 | 2 | 0.68 | 0.06 | 0.58 | 0.80 |
| 1300 | 38 | 2 | 0.65 | 0.06 | 0.54 | 0.77 |
| 1400 | 36 | 1 | 0.63 | 0.06 | 0.52 | 0.76 |
| 1500 | 21 | 1 | 0.60 | 0.06 | 0.48 | 0.74 |
| 1700 | 20 | 2 | 0.54 | 0.07 | 0.42 | 0.70 |

**B. Foster birds**

| time | n. risk | n. event | survival | se | lower 95% CI | upper 95% CI |
|---|---|---|---|---|---|---|
| 100 | 38 | 4 | 0.90 | 0.05 | 0.80 | 1.00 |
| 500 | 34 | 2 | 0.84 | 0.06 | 0.73 | 0.97 |
| 600 | 20 | 1 | 0.80 | 0.07 | 0.68 | 0.95 |
| 800 | 16 | 1 | 0.75 | 0.08 | 0.61 | 0.93 |
| 1000 | 15 | 1 | 0.70 | 0.09 | 0.54 | 0.90 |
| 1400 | 14 | 1 | 0.65 | 0.10 | 0.49 | 0.87 |
| 1500 | 11 | 1 | 0.59 | 0.10 | 0.42 | 0.84 |
| 1700 | 10 | 1 | 0.53 | 0.11 | 0.36 | 0.80 |
